# Decellularisation and characterisation of porcine pleura for lung tissue engineering

**DOI:** 10.1101/2023.06.20.545830

**Authors:** Trisha Vikranth, Tina Dale, Nicholas R Forsyth

## Abstract

Decellularisation offers a broad range of biomimetic scaffolds of allogeneic and xenogeneic origins, exhibiting innate tissue-specific characteristics. We explored a physico-chemical method for decellularising porcine pleural membranes (PPM) as potential tissue-engineered surrogates for lung tissue repair. Decellularised PPM (dPPM) was characterised with histology, quantitative assays, mechanical testing, and sterility evaluation. Cytotoxicity and recellularisation assays assessed the biocompatibility of dPPM. Haematoxylin and Eosin staining showed an evident reduction in nuclei in dPPM, confirmed with nuclear staining and analysis (****p < 0.0001). Sulphated glycosaminoglycans (sGAG) and collagen histology demonstrated minimal disruption to the structural assembly of the core extracellular matrix (ECM) in dPPM. Confocal imaging demonstrated realignment of ECM fibres in dPPM against native control. Quantitative analysis defined a significant change in the angular distribution (****p < 0.0001) and coherence of fibre orientations (***p < 0.001) in dPPM versus native ECM. DNA quantification indicated ≥ 85% reduction in native nuclear dsDNA in dPPM (**p < 0.001). Collagen and sGAG quantification indicated reductions of both (**p < 0.001). dPPM displayed increased membrane thickness (*p < 0.05). However, the Youngs modulus (447.8 ± 41.9 kPa) and ultimate tensile strength (5080 ± 2034.5 kPa) of dPPM were comparable with that of native controls at (411.3 ± 8.1 kPa) and (3933.3 ± 1734.5), respectively. *In vitro* cytotoxicity and scaffold biocompatibility assays demonstrated robust human mesothelial cell line (MeT-5A) attachment and viability. Here, we define a decellularisation protocol for porcine pleura that represents a step forward in their potential application as bioscaffolds in lung tissue engineering.

**Impact statement:** Design and development of ‘off the shelf’ tissue engineered products can rely on decellularisation of native tissue that can be functionalised with cell ingress from surrounding host microenvironment. We describe a reproducible decellularisation method for porcine pleural membranes. Histology, mechanical testing, and biocompatibility studies demonstrated protocol efficiency in adequate removal of native tissue cellularity and retention of gross microarchitecture and bioactivity in the decellularised pleura. The study represents a step forward in establishing an off the shelf potential of decellularised pleura as site-specific mechanical barriers in curtailing prolonged air leaks and promoting spontaneous tissue regeneration with relevant physiological cues.

## Introduction

Advances in tissue engineering (TE) and biomaterial sciences have revolutionised the scope of biomimetic transplant fabrication, capable of replacing and restoring damaged tissue structure and function. The classic paradigm in engineering a tissue construct is in the use of a mechanically and biochemically conditioned scaffold that modulates cell proliferation, differentiation, and constructive tissue remodelling. Essentially, recapitulating the structural and functional attributes of native ECM. Optimised whole organ and membrane decellularisation protocols have produced biological scaffolds that exhibit the structural and functional attributes of native ECM. With ECM composition and architecture being relatively conserved amongst species^1,2^, the options of using accessible, economically viable and biocompatible xenogeneic scaffolds have considerably expanded with decellularisation. Decellularised porcine small intestinal submucosa (SIS) ^3^ and urinary bladder matrix (UBM)^4^, derived by mechanical delamination of the tunics have shown excellent retention of native ECM histoarchitecture and biomechanics, gaining clinical applications in genitourinary reconstruction ^5–7^ and soft tissue augmentation ^8–11^. Decellularised porcine dermis processed as freeze-dried sheets, has been used clinically to promote reepithelialisation in full- and partial thickness wounds and burns ^12,13^.

ECM is a complex three-dimensional (3D) biomolecular assembly of structural and functional proteins and proteoglycans organised in a tissue-specific pattern. In addition to providing structural framework, ECM is equipped with functional machinery that establishes a dynamic two-way communication between cells and their external microenvironment ^14,15^. Characterisation studies have defined the ECM core to be composed of fibrous proteins predominantly, collagens, elastin, fibronectin and laminins embedded within a lubricating and cushioning hydrogel, rich in proteoglycans (PG) ^14,16,17^. Collagens and elastin generate the mechanical resilience and topographical cues necessary for maintaining tissue structure and function. Fibronectin and laminins facilitate cell adhesion, migration, and the release of growth factors as physiological response to tissue damage and injury. Proteoglycans (PG) with attached glycosaminoglycan units provide hydration, compressive strength and a repository of secretory modulators that include vascular endothelial growth factor (VEGF), fibroblast growth factor (FGF), epidermal growth factor (EGF) and transforming growth factor - beta (TGF-β) ^18^.

Pulmonary pleurae are the thin membranous lining that cover lungs and the thoracic cavity. Microscopically, they are defined by a superficial mesothelial monolayer resting on a connective tissue matrix rich in collagens and elastin ^19,20^. Pleurae form the protective low friction barrier aiding smooth movement of lungs within the thoracic cavity during breathing. The mesothelium secretes a diverse profile of soluble factors that facilitate immunomodulatory and inflammatory responses to pleural injury, exhibiting innate ability in wound healing and tissue regeneration. However, the regenerative potential is often limited to the extent and severity of pleural injury and disease ^21^.

Persistent air leaks following routine lung surgery that last longer than 5 days are defined as PAL. At an incidence of 23%, PAL are a source of significant post-operative morbidity associated with extended chest tube durations and secondary pulmonary complications. ^22,23^. Current surgical practice involve intra-operative closures such as buttressing staple lines, pleurodesis, and use of biological and composite sealants ^24,25^. Fibrin, a routinely used natural sealant known for its biocompatibility, lacks mechanical strength in sustaining long-term closure of surgical incisions. Retrospective studies of air leak management are yet to provide compelling evidence to suggest a superior technique in preventing PAL. Most studies report a marginal decrease in the incidence of PAL with no significant reduction in chest tube durations or hospital admissions ^23,26–29^. On the contrary, use of sclerosing agents and synthetic sealants are associated with acute inflammatory responses in patients.

We aimed to develop a reproducible decellularisation protocol for porcine pleura and assess protocol efficiency in adequate cell removal and preservation of structural and functional characteristics of pleural ECM. We characterised dPPM scaffolds for histological features, mechanical behaviour, and biocompatibility to provide proof of concept for their putative application as a TE acellular graft in lung tissue repair.

## Materials and Methods

### Decellularisation of PPM

Fresh pig lung sourced from a local abattoir was used for pleural membrane excision. Excision was carried out in a class I dissection hood with lungs placed in crushed ice. PPM were peeled gently and transferred to petri-dishes containing chilled phosphate buffered saline (PBS, 1x without calcium and magnesium, pH 7.4 ± 0.1, Corning®, USA). Excised membranes were rinsed thoroughly in deionised water to remove blood and cell debris.

As in figure 1, PPM disruption and cell lysis was initiated with 5X freeze-thaw cycles of snap freezing at −80°C for 15-20 min followed by thawing at room temperature for 25-40 min. Repeated washing with deionised water ensured removal of extraneous matter and cell debris. Samples were next immersed in a decellularisation buffer, (1% (v/v) Triton-X 100, 0.5% sodium deoxycholate in 10 mM Tris buffer pH 7.6 (both Sigma – Aldrich) and 30 µg/mL DNase I solution (Invitrogen™, USA), for 48 hours at 37°C under mild agitation. Treated samples were refrigerated and washed 6X with deionised water at 8 hour intervals and then 6X with PBS at 12 hour intervals. dPPM were stored at 4°C in PBS containing 1X penicillin/streptomycin/amphotericin B solution (100X PSA containing 10,000 units potassium penicillin/mL, 10,000 µg streptomycin sulphate/mL and 25µg amphotericin B/ml in 0.85% saline, Lonza, Switzerland) for two weeks before being transferred to −80°C, for long term preservation. Three biological replicates with five technical repeats for native and dPPM samples were studied.

**Figure 1.**
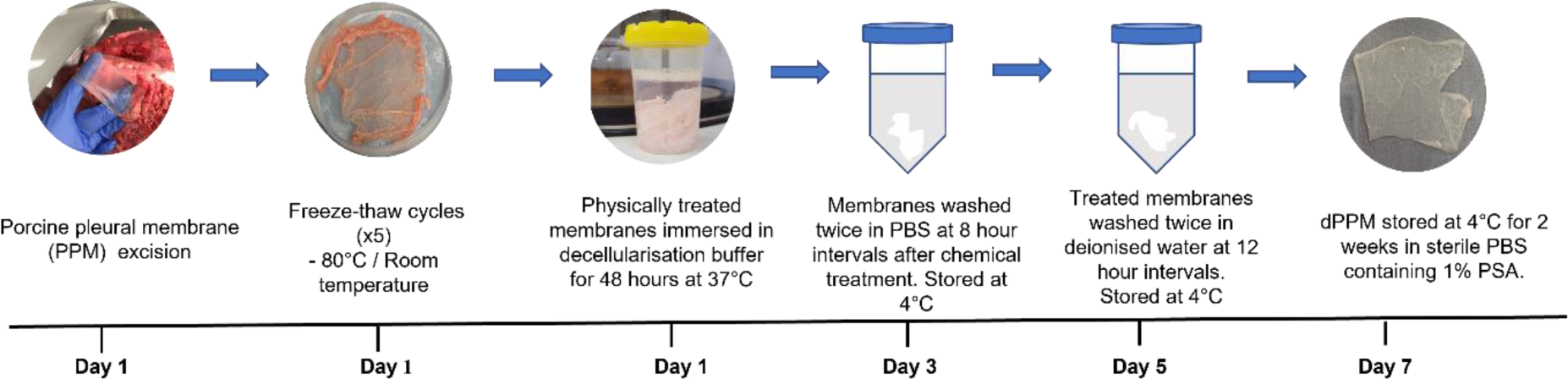
Schematic representing protocol for porcine pleural membrane (PPM) decellularisation.

### Histology

dPPM was stained with H&E (Hematoxylin solution, Gill no.2, Sigma Aldrich, aqueous Eosin Y, Thermo scientific) to assess cellular content and histoarchitecture. Picrosirius red (PSR) (Sigma – Aldrich) staining was used to visualise collagen distribution and orientation while Alcian blue (1% Alcian blue in 3% acetic acid, Thermo Fisher scientific, MA, USA) staining was used to qualitatively examine sGAG content. Elastin fibres were examined using standard IHC protocol for Verhoeff-Van Gieson (VVG) staining. Stained samples were dehydrated using a graded alcohol series and mounted with DPX (360294H, Analar) mounting media. Phase contrast and bright field images were acquired at 20X magnification (EVOS digital imaging systems, AMG Washington).

### Confocal imaging and analysis

Fresh native and dPPM samples were imaged under a laser scanning confocal microscope (Olympus Fluoview FV1200, Japan) at 20X magnification. Autofluorescence of ECM fibres were detected in the samples when excited with a 488 nm multiline Argon laser. Image acquisition and control was carried out using the integrated FV10 – ASW software. The 2D images acquired, were in ImageJ compatible OIB (Olympus image binary) formats with a resolution of 1024 x 1024 pixels (physical size = 636 microns x 636 microns, 16-bit).

OrientationJ plugin for ImageJ/Fiji was used for image analysis. A 10-pixel Gaussian analysis window with an estimated cubic spline gradient, was specified for the distribution and measurement modes in OrientationJ. Structure tensors computed by sliding the Gaussian analysis window across the image, was used for determining local orientations of ECM fibres. Orientations ranging from – 90° to + 90° from the horizontal, were represented as colour-coded orientation maps and histograms of the distribution of fibre orientations in native and dPPM, using the OrientationJ distribution mode. Measurement mode of OrientationJ quantified differences in coherency and orientation of ECM fibres between native ECM and dPPM. Each image was analysed with 8 regions of interest (ROI) selected using the rectangular select tool in ImageJ. Best fitting ellipses created for each ROI defined features of the gradient structure tensors with respect to fibre orientations and coherency. Angular distribution of fibres and the degree to which these features were oriented in each ROI, was tabulated as orientation and coherency. Orientation values were represented in degrees and coherency values as an index between 0 and 1. Coherency of 1 indicated a single dominant fibre orientation in the ROI while 0 indicated fibre isotropy in orientation.

### Nuclear labelling and DNA quantification

Nuclear DNA integrity and gross cell morphology in dPPM was assessed with 4’,6’- diamidino - 2 - phenyl indole (DAPI, Sigma-Aldrich) staining and fluorescence microscopy (Nikon Eclipse Ti, Japan). Nuclear quantitation was performed using the analysis plugin of ImageJ (Rasband, W.S., ImageJ, U.S. National institutes of Health, USA).

Native PPM and dPPM samples were desiccated in a hot air oven overnight to a constant weight (approximately 25 mg. dry weight) prior to DNA quantification. Samples were treated overnight with proteinase K (1.25 mg/mL, Sigma-Aldrich) at 55°C, to obtain clear membrane digests. Residual dsDNA in dPPM and native digests was quantified using the Quant-iT^™^ PicoGreen^™^ dsDNA assay kit (Thermo Fisher Scientific, USA) following manufacturer’s instructions. Samples were read at 480 nm and 520 nm excitation and emission wavelengths, respectively, on a fluorescence microplate reader (Biotek Synergy 2 plate reader, USA). dsDNA content was calculated using the standard calibration curve, following manufacturer’s instructions, and expressed as µg/mg. dry weight of tissue.

### sGAG quantification

Samples were first dried and digested as described above. Quantification of sGAG content was established using the 1,9-dimethyl methylene blue assay (DMMB, abcam, UK) following manufacturer’s instructions. Digested samples (50 µL) were aliquoted in a 96-well microplate. 200 µL of the DMMB solution (16 mg DMMB, 0.76 glycine, 0.595 g NaCl, 23.75 mL of 0.1 M HCl and distilled water to 250 mL) were added to each well using an automated dispenser. Sample absorbance was immediately measured at 530 nm on a Synergy 2 microplate reader. sGAG content expressed as µg/mg. dry weight of tissue, was determined against a standard calibration curve, following manufacturer’s instructions.

### Collagen quantification

Samples were prepared using acid-pepsin extraction guidelines from the manufacturer (Sircol™, insoluble collagen assay S2000, Biocolor Ltd.). Samples were then dried and treated overnight with the Acid-pepsin solution (0.1 mg/mL pepsin in 0.5 M acetic acid, Sigma – Aldrich) at 4°C on a mechanical shaker. The digested samples were centrifuged at 3000 g for 10 minutes to obtain the residual tissue for the insoluble Sircol binding collagen assay following manufacturer’s guidelines. Sample absorbance was measured at 556 nm on a Synergy 2 microplate reader. Insoluble or native collagen content was determined against a standard calibration curve and expressed as µg/mg. dry weight of tissue.

### Biomechanical studies

Membrane thickness of native and dPPM samples were measured using an optical tensiometer (Attention, Biolin Scientific, Manchester, UK) and ImageJ software. Fresh dPPM and native samples cut into 4 cm x 1cm sections, were clamped between the upper (mobile) and lower (stationary) grips of a benchtop tensile testing apparatus (Bose ELF 3200, Minnetonka, USA). Samples were subjected to uniaxial tension to failure, at a load of 22N and relative axial displacement of 10 mm at 0.1mm/sec. Initial length (L_0_) was kept consistent at 1 mm using a digital calliper. WinTest®7 software was used for data acquisition. Resulting stress-strain curves were used to calculate Youngs modulus and ultimate tensile strength of samples.

### Sterility evaluation

For sterilisation with UV, dPPM samples (n = 5) were irradiated using the pre-programmed sterilisation setting (maximum wavelength of 253.7 nm, 90 seconds exposure) on the GS Gene Linker UV chamber (Bio-Rad laboratories, California, USA). Samples placed in petri dishes received three repeats of irradiation for top and bottom surfaces of the dish. Alternatively, dPPM samples (n = 5 for each treatment) were treated with either 10X PSA, 90% industrial methylated spirits (IMS), or 0.1% peracetic acid (PAA) in 4% aq. Ethanol for 30 min. After treatment, samples were rinsed in sterile PBS and transferred to 24-well plates. Dulbecco’s modified eagles medium (DMEM, 4.5 g/L glucose, sodium pyruvate without L-glutamine, Corning), containing 10% fetal bovine serum (FBS, Biosera), was added to each well and the plates incubated under standard conditions (37°C, 5% CO_2_ and 95% relative humidity). Sample wells were assessed visually at day 0, 3 and 7 for infection, to determine sterilisation efficiency.

### *In vitro* cytotoxicity assay

Human mesothelial cells (MeT-5A, ATCC CRL-9444™, Virginia, USA) were cultured in cFAD medium ^30^, composed of DMEM and Hams F-12 (Lonza), (3:1, v/v), supplemented with 10% FBS, 1% non-essential amino acids (Lonza), 2mM L-glutamine (200mM, Lonza), 1% sodium pyruvate (100 mM, Lonza), 1% PSA, 24 µg/mL Adenine (Sigma - Aldrich), 0.4 µg/mL hydrocortisone (Sigma - Aldrich), 0.13 µg/mL triiodothyronine (Sigma - Aldrich), 10 ng/mL epidermal growth factor (EGF, Sigma - Aldrich), 5 µg/mL transferrin (Sigma - Aldrich), 5 µg/mL insulin (Sigma - Aldrich). Cells were seeded at 0.1 x 10^5^ cells/mL on collagen coated 24-well plates and incubated in standard cell culture conditions. After 48-hours, 5 mm x 5 mm sections of sterilised dPPM were placed in the wells. Cells incubated in cFAD with no contact with dPPM and cells treated with 70% methanol for 30 min served as controls for live-dead staining, respectively. Fluorescence viability assay using the Live/Dead staining kit (Invitrogen™ L3224) was carried out following manufacturer’s instructions. Sample wells were imaged using dual channel filters, FITC for live cells and Texas Red for dead cells. Cell viability was determined using Trypan blue (Lonza) exclusion assay to estimate viable cell counts at specific timepoints from day 0 – day 5 of incubation.

### Scaffold biocompatibility

Sterilised dPPM cut into 5 mm x 5 mm sections, were mounted on sterile 24-well CellCrown™ inserts (Scaffdex Oy, Sigma-Aldrich, USA) and placed in wells of a 24-well plate. MeT-5A cells were labelled with a live cell tracking dye (Vybrant® DiO (1mM), Invitrogen™ V22889) following manufacturer’s recommended protocol and seeded at a density of 0.3 x 10^5^ cells/cm2, on dPPM inserts. Sample wells were incubated under standard cell culture conditions and imaged periodically. Labelled MeT-5A cells seeded on tissue culture plastic (TCP) and sterile dPPM inserts served as positive and negative controls, respectively.

### Statistical analysis

Statistical analysis was performed using GraphPad Prism version 9.0 software (GraphPad Software Inc., San Diego, CA). Data was represented as Mean ± standard deviation (SD) at 95% confidence intervals for each study. Results for dPPM and native samples from mechanical testing and quantitative assays were compared using an unpaired students t-test. (*p < 0.05, **p < 0.001, ****p < 0.0001) was considered statistically significant.

## Results

### PPM decellularisation and histology

Macroscopically, native PPM (Fig 2A) appeared white and opaque following decellularisation (Fig 2B). Fluorescence imaging of DAPI stained nuclei in native (Fig 2C) and dPPM (Fig 2D) showed a marked reduction in cellularity in dPPM.

**Figure 2.**
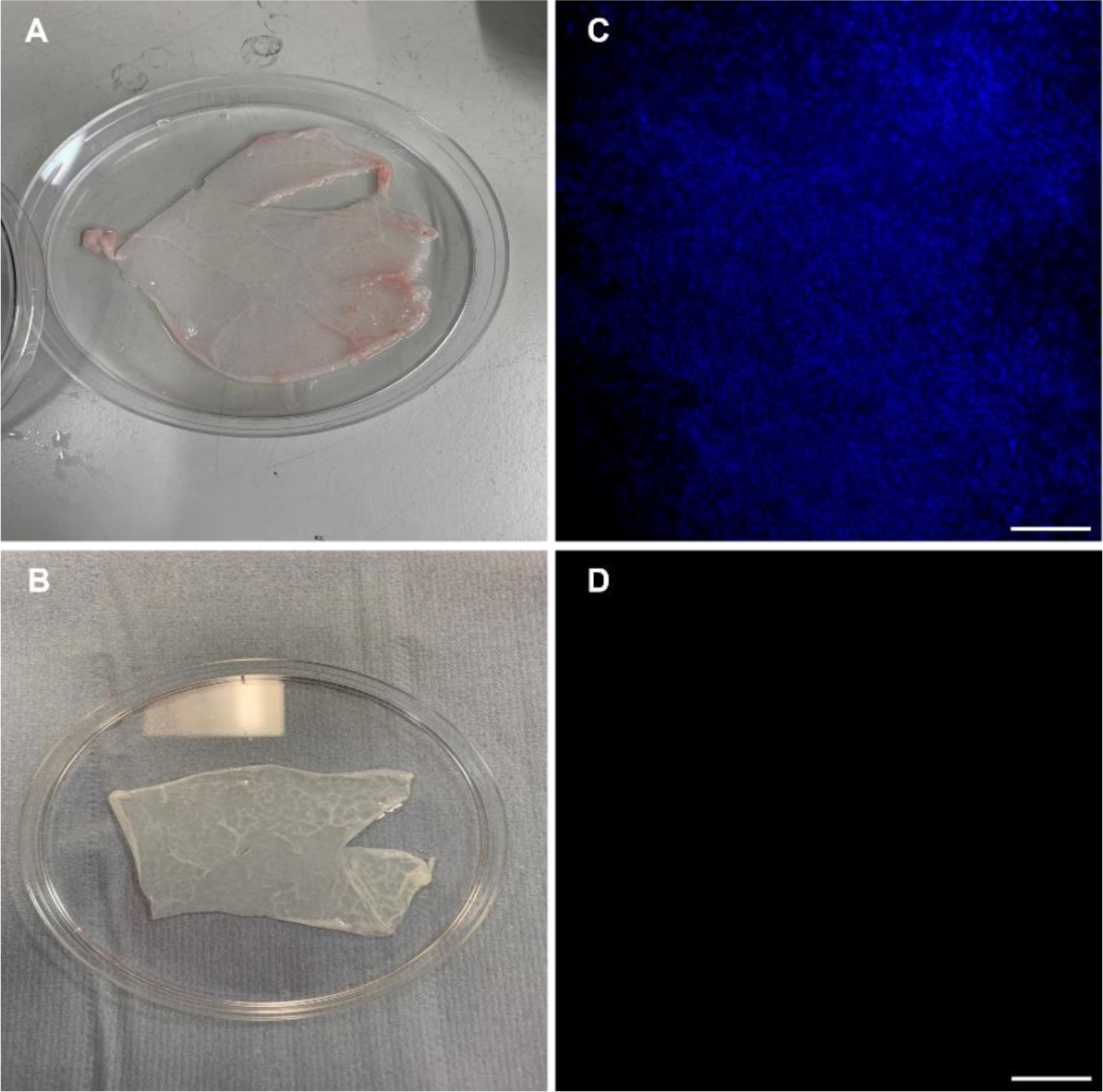
Characterisation of native and decellularised PPM. (A,B) represent macroscopic changes to the native PPM (A) following decellularisation (B). (C,D) represent the visible reduction in DAPI stained nuclei in dPPM (D) against native PPM (C). Scale bar = 200 µm.

Histological features in dPPM demonstrated changes relative to the microanatomy of native PPM (Fig 3A), with an overall reduction in nuclei (Fig 3B). Similarly, Alcian blue staining suggested a reduction in native sGAG content (Fig 3C) in dPPM (Fig 3D). PSR labelling under cross-polarised light in native (inset Fig 3E) and dPPM (inset Fig 3F), demonstrated reduced collagen staining in dPPM in comparison. The gross organisational structure of collagen in native tissue (Fig 3E) under polarised light remained relatively conserved in dPPM (Fig 3F). VVG staining for elastin fibres (black) in native (Fig 3G) and dPPM (Fig 3H) also demonstrated reduction in staining intensity for elastin following decellularisation.

**Figure 3.**
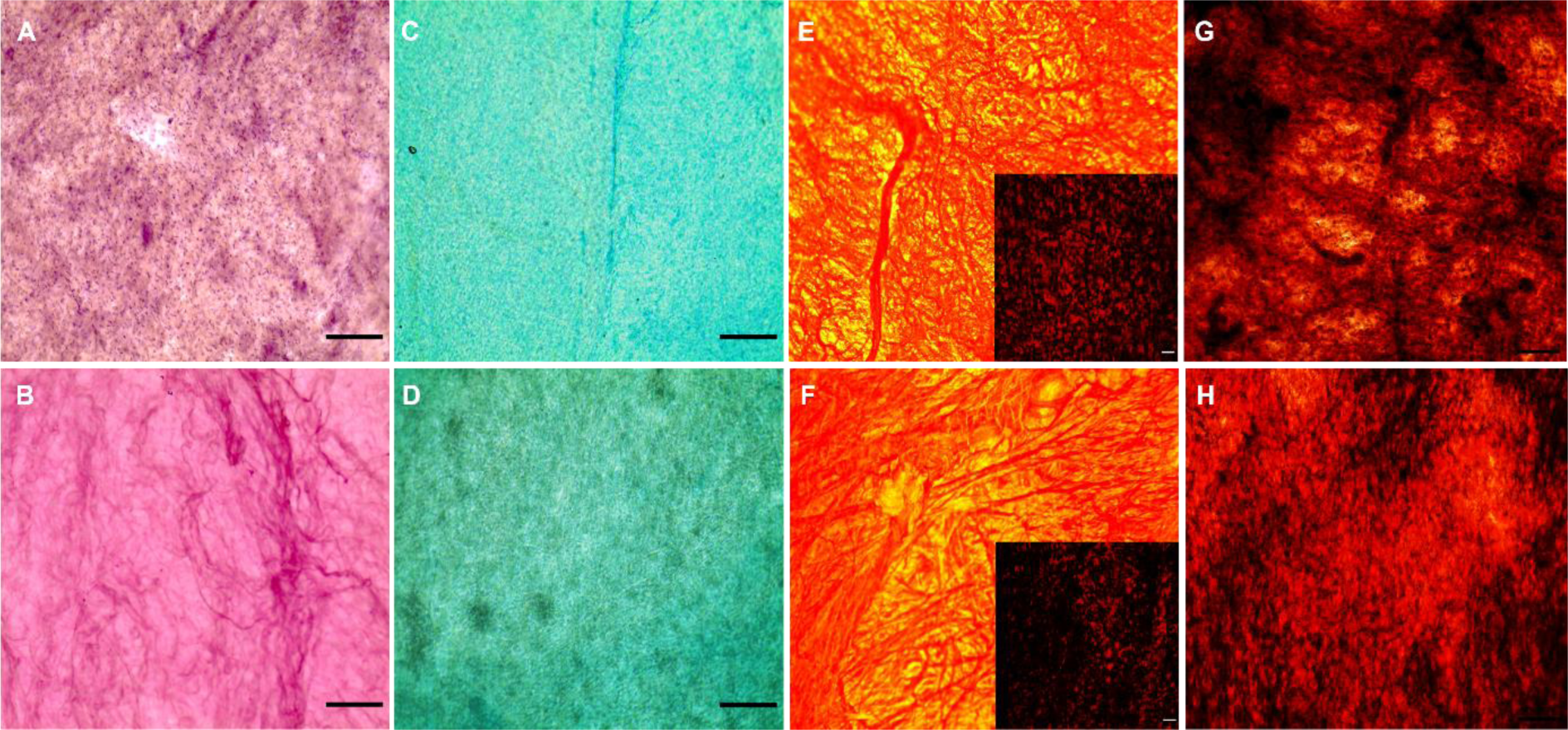
Histological features of native and decellularised PPM. (A,B) Hematoxylin and Eosin (H&E) of native (A) and dPPM (B). (C,D) Alcian blue staining in native PPM (C) and dPPM (D). (E,F) Picrosirius red staining in native (E, inset under cross-polarised light) and dPPM (F, inset under cross-polarised light). (G,H) Verhoeff-Van Geison (VVG) staining in native (G) and dPPM (H). Scale = 100 µm

### Confocal imaging and analysis

Microstructural changes in the fibrous organisation of native ECM after decellularisation was quantified using the OrientationJ plugin of ImageJ. Images of fresh native (Fig 4A) and dPPM (Fig 4B) captured with confocal microscopy, were analysed for orientation and coherency. Colour-coded orientation maps generated for native (Fig 4C) and dPPM (Fig 4D) highlighted predominant orientations of fibres for each, which on further analysis generated a distinct peak for the distribution of orientations of dPPM fibres in contrast with a plateau – like multimodal distribution of fibre orientations in native (Fig 4E). Quantitative measurements for orientation also indicated a significant increase in coherency of fibre orientations for dPPM (Fig 4F) against native (***p < 0.001). Degree of orientation (Fig 4G) was in concurrence with Fig 4E, where native fibres demonstrated a predominant angular position of −5° in contrast with the dPPM fibres assuming an alignment at −44° from the horizontal. (****p < 0.0001).

**Figure 4.**
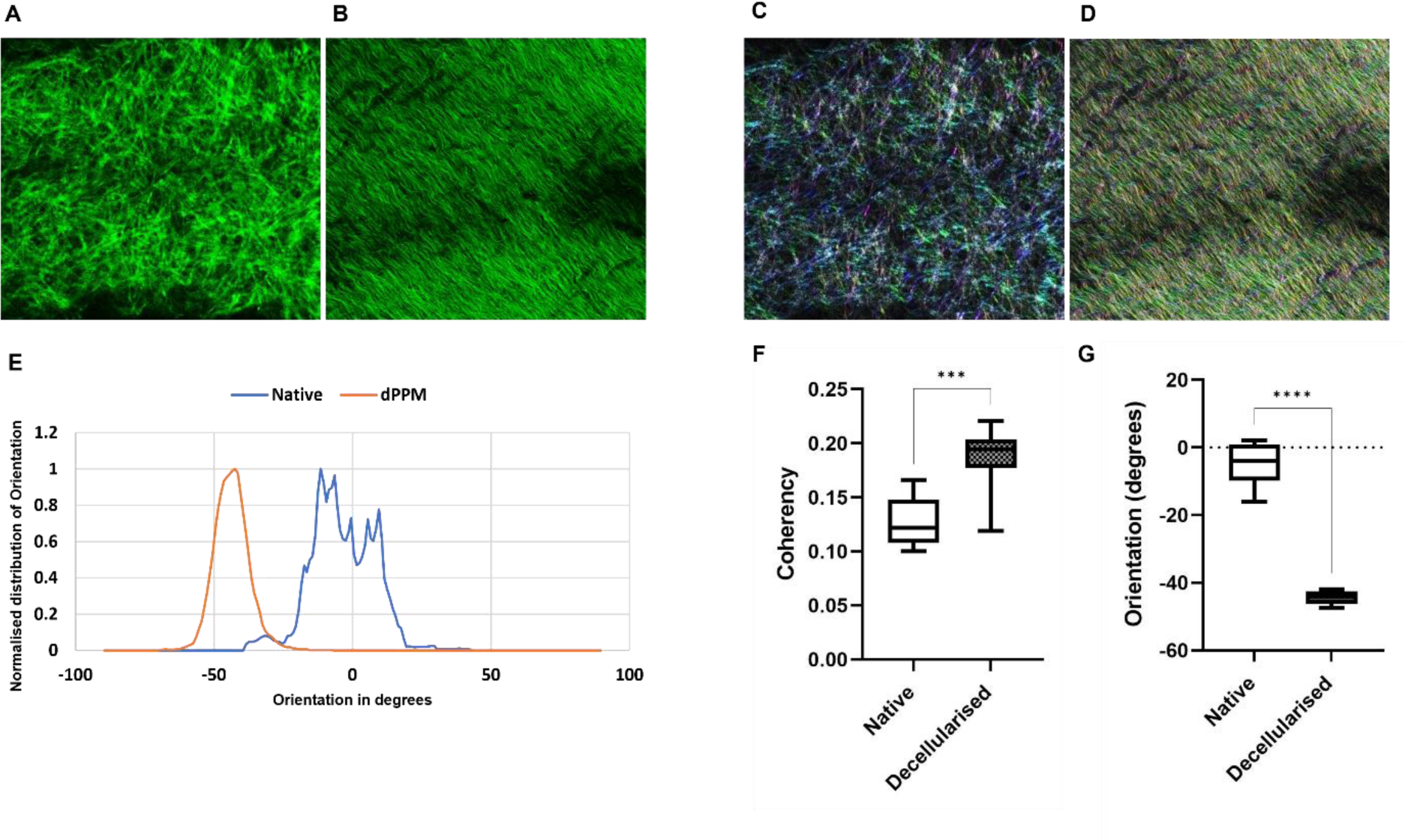
Microstructural alignment of ECM fibres in native and dPPM. Native (A) and dPPM (B) ECM observed under confocal microscopy. Scale = 200 µm. (C,D) represents colour-coded fibre orientation maps of native (C) and dPPM (D) samples. (E) represents angular distribution of ECM fibres in native and dPPM. (F) represents coherency in fibre orientations in native and dPPM. (G) represents dominant ECM fibre orientations in native and dPPM. (n = 3, ***p < 0.001, ****p < 0.0001, mean ± SD).

### Bioquantitative assays

Quantification of DAPI stained nuclei (Fig 5A) in native (2.1 x 10^5^ ± 0.3 x 10^5^ nuclei/cm^2^) and dPPM (0.05 x 10^5^ ± 0.06 x 10^5^ nuclei/cm^2^) indicated a 40-fold reduction in dPPM vs. native (n = 5, ****p < 0.0001). This was reflected in overall DNA content (Fig 5B) in dPPM (15.2 ± 5.9 ng/mg. of tissue) versus native tissue (123.5 ± 74.6 ng/mg. of tissue), demonstrating a significant reduction of over 85% of native nuclear DNA with decellularisation (n = 5, **p < 0.01). sGAG quantification (Fig 5C) of dPPM (4.2 ± 0.4 µg/mg) versus native (7.9 ± 1.6 µg/mg of tissue) revealed a 53.5% reduction in content following decellularisation (n = 5,**p < 0.01). Insoluble collagen content (Fig 5D) was reduced in dPPM (52.7 ± 8.4 µg/mg of tissue) versus native control (76.0 ± 10.5 µg/mg of tissue) with an overall 31% reduction (n = 5,**p < 0.01).

**Figure 5.**
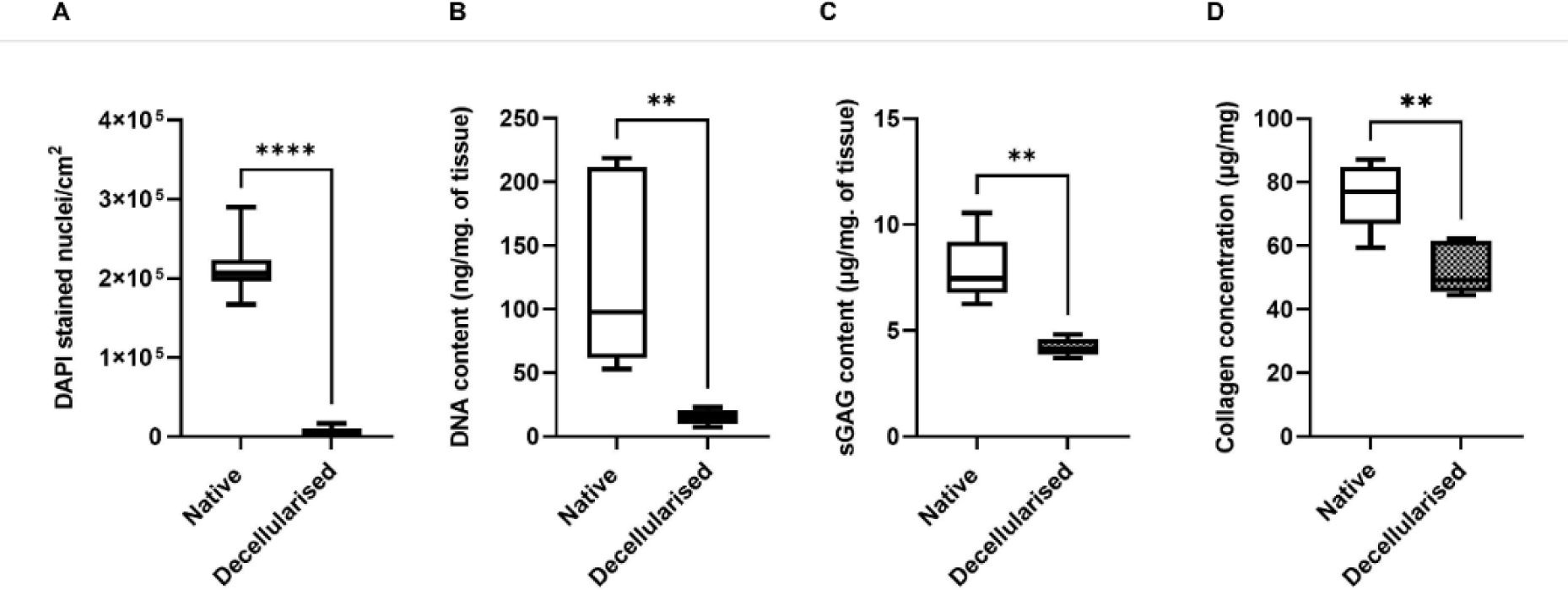
Protocol efficiency in PPM decellularisation. (A) DAPI stained nuclei in native vs. dPPM. (B) DNA content in native vs. dPPM. (C) sGAG content in native vs. dPPM. (D) Collagen content in native vs. dPPM. (n=5, **p < 0.01, ****p ≤ 0.0001). Data represented as Mean ± SD.

### Sterility assessment

UV and 10% PSA treated samples displayed turbidity and colour change in culture media, in all wells by day 3 of incubation. Similar observations were made with IMS treated samples with 33% of the wells showing infection by day 3 (Fig 6A). 0.1% PAA treated samples demonstrated no change to medium opacity and colour throughout the incubation period (Fig 6A,B).

**Figure 6.**
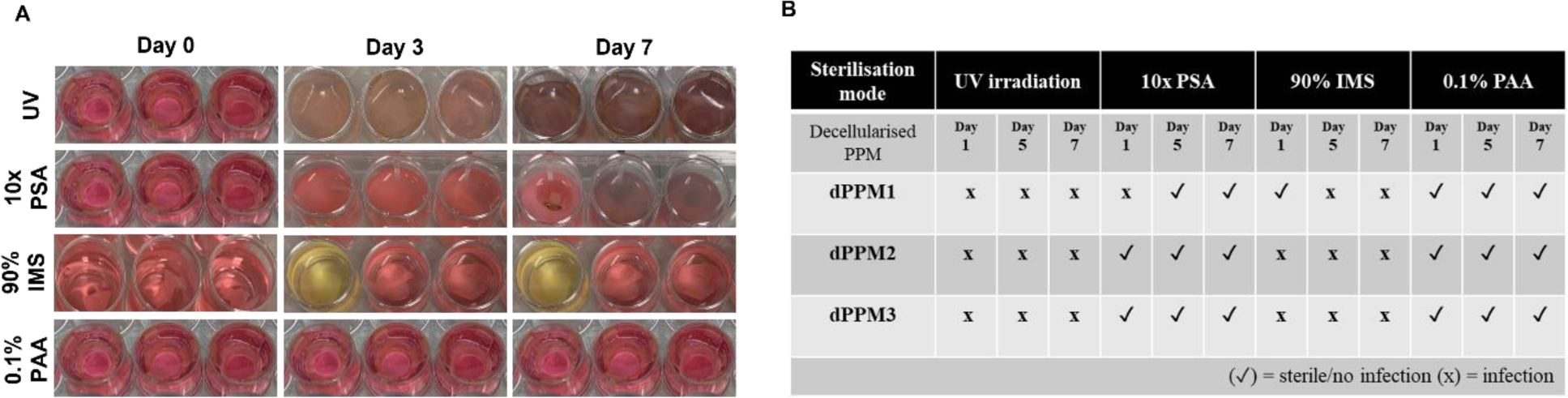
Sterility assessment of dPPM. (A) Sterilisation efficiency of UV, 10X PSA, 90% IMS and 0.1% PAA at day 0 - day 7 (B) dPPM sterility logs from day 0 - day 7 (n = 3, for each treatment)

### Mechanical characterisation

dPPM displayed a 32.6% increase in membrane thickness (194.5 ± 11.3 μm) against native (147.2 ± 27.2 μm) (n = 3, *p < 0.05) (Table 1). Stress-strain curves of dPPM under uniaxial tension demonstrated viscoelastic characteristics with an initial toe region, followed by the linear and failure regions of response. Core mechanical characteristics of native PPM with an estimated Youngs modulus of 411.3 ± 8.1 kPa and ultimate tensile strength of 3933.3 ± 1734.5 kPa was conserved in the dPPM demonstrating comparable Youngs modulus (447.8 ± 41.9 kPa) and ultimate tensile strength (5080 ± 2034.5 kPa). 0.1% PAA sterilised dPPM retained comparable mechanical strength and behaviour against native control with an estimated Youngs modulus of 429.8 ± 34.5 kPa and ultimate tensile strength of 5413.3 ± 1510.1 kPa (Table 1).

**Table 1.** Biomechanical characterisation of native and decellularised PPM

| Mechanical properties of PPM under uniaxial tension |  |  |  |
| --- | --- | --- | --- |
| | Mean membrane thickness ($\mu\text{m}$) | Mean Youngs modulus (kPa) | Mean Ultimate tensile strength (kPa) |
| Native PPM | $147.23 \pm 27.15$ | $411.25 \pm 8.12$ | $3933.33 \pm 1734.51$ |
| Decellularised PPM | $194.53 \pm 11.30^*$ | $447.80 \pm 41.85$ | $5080 \pm 2034.5$ |
| 0.1% PAA sterilised dPPM | $198.22 \pm 9.21^*$ | $429.27 \pm 34.51$ | $5413.33 \pm 1510.14$ |

Data represented as Mean ± SD, $n = 3$ . $*p < 0.05$ vs native PPM

### Contact cytotoxicity assay

Live/Dead staining of MeT-5A cells in contact with sterilised dPPM on days 1, 3 and 5 demonstrated negligible presence of homodimer (EthD-1) stained dead cells (Fig 7A). Trypan Blue exclusion assay provided quantitative evidence of negligible cytotoxicity with the average number of viable cells/cm^2^ to dead cells/cm^2^ being significant from day 0 - day 5 (p < 0.05) (Fig 7B).

**Figure 7.**
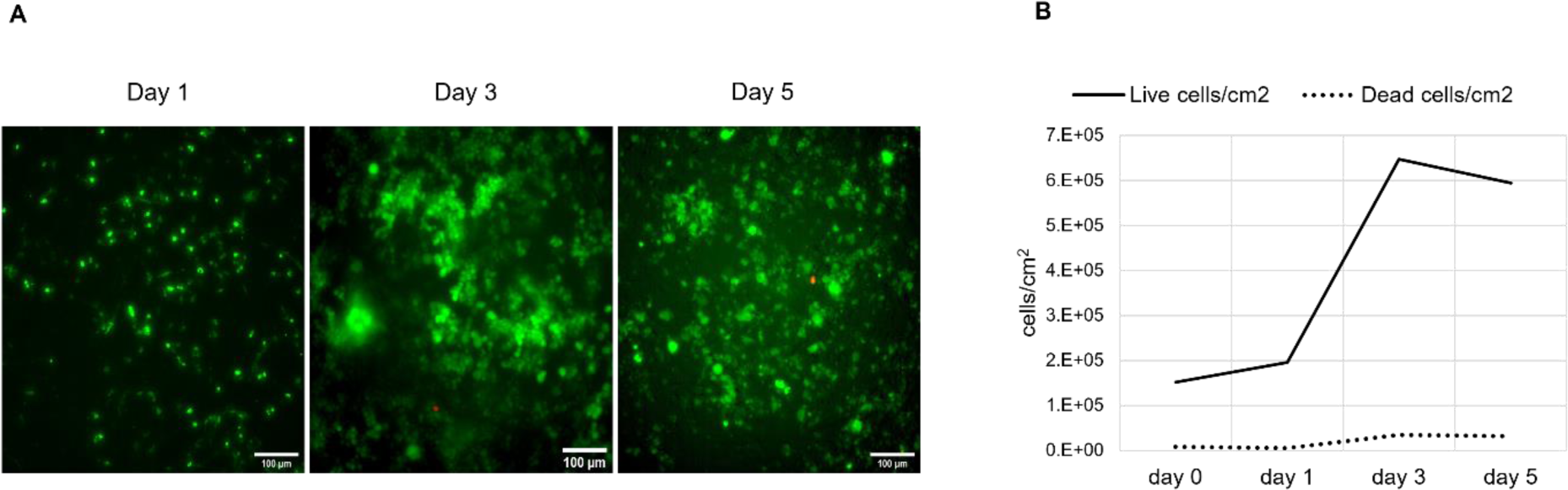
Biocompatibility of decellularised PPM with contact cytotoxicity assay. (A) Live (green) / Dead (red) staining of seeded MeT-5A cells in contact with sterilised dPPM on day 1 – day 5. (B) Viability of MeT-5A cells in contact with sterilised dPPM from day 0 – day 5. (n =3, *p ≤ 0.05). Data represented as Mean ± SD.

### Scaffold biocompatibility

MeT-5A seeded - dPPM indicated cellular attachment on day 1. Scaffolds were aseptically transferred to fresh wells for imaging, to confirm attachment of cells to the scaffold and not the culture plate. Morphology and proliferation of MeT-5A cells seeded on dPPM resembled the control MeT-5A cells seeded directly on TCP (Fig 8). Lipophilic carbocyanine dye used to label MeT-5A permeates the cell membrane and laterally diffuses to stain the entire cell surface. With proliferation of cells in long-term culture, the dye is diluted over time, giving cells a granular appearance as seen in both, the seeded control and dPPM wells, on day 9 and day 15 ^31^ (Fig 8).

**Figure 8.**
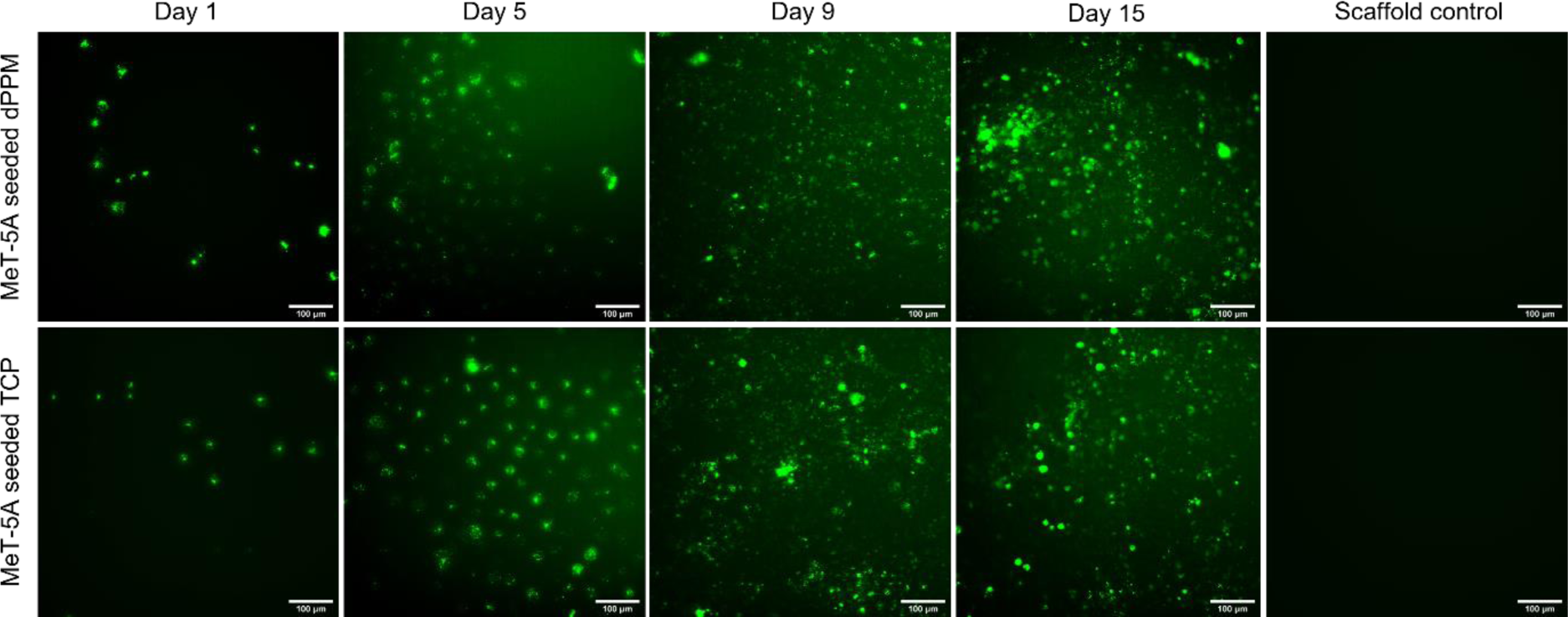
Recellularisation of sterilised dPPM using labelled MeT-5A cells. Attachment and proliferation of MeT-5A cells labelled with a live cell tracker, was observed from day 1 through day 15, as represented. MeT-5A cells seeded on standard TCP and unseeded dPPM (scaffold only) served as positive and negative controls, respectively.

## Discussion

Pioneered by Meezan et al in 1975 ^32^, whole organ and membrane decellularisation have gained a strong foothold in scaffold fabrication approaches in tissue engineering. Preservation of tissue-specific microarchitecture, biomolecular makeup, and mechanical attributes of native ECM with reduced risk of immunogenicity are the hallmarks of an optimal decellularisation process. Porcine biological scaffolds as human-scale substitutes, owing to the evolutionarily conserved and comparable ECM characteristics, have found clinical applicability in cardiovascular, ^33–35^ cranial, ^36,37^ and bladder ^3,38,39^ reconstruction.

Even with advanced minimally invasive surgical techniques and intraoperative wound closure measures, approximately 5% - 10% of patients suffer from post-surgical PAL lasting longer than 7 days, often requiring surgical intervention ^23,40–42^. Kanzaki et al ^43^ in 2008, described a TE alternative for functional closure of compromised pleura using temperature responsive smart polymers to develop transplantable autologous dermal fibroblast cell sheets. Irrespective of outcome, time-bound logistics in deriving and preserving viable autologous cell-sheets, transportation and storage are potential translation barriers in the clinical application.

We hypothesised the potential of decellularised porcine pleura as site-specific mechanical barriers or adjuncts, to reinforce current closure techniques in reducing the incidence and severity of PAL. Although considerable studies support wound closure techniques that include surgical stapling^44^, pleurodesis^45^ and application of biological and synthetic sealants^46^, the incidence and severity of PAL associated complications, seen with spontaneous and traumatic penumothoraces pose a significant clinical challenge in air leak management. With this in mind, our study optimised a protocol to derive transplantable acellular pleural scaffolds for potential tissue-specific applications in pleural repair.

Studies have adopted various physical, chemical, enzymatic, and combinative methods to optimise decellularisation, specific to the source and nature of the tissue, composition, density, and intended clinical application ^17,47–49^. The two-step decellularisation approach in our study initiated with freeze-thaw cycles and treatment with a detergent-nuclease buffer system, was in consideration of preserving the viscoelastic nature of the native pleura. Intracellular ice crystal formation during freeze-thaw cycles induce cell lysis and subsequent removal of cell debris with periodic washes ^47,48^. Tissues including peripheral nerve grafts, heart valve leaflets, and dermis have been decellularised with this technique, characterised by minimal disruption in gross ECM morphology. However, considerable retention of nuclear material was reported with this process alone ^50–52^. Our subsequent treatment with a detergent buffer containing low concentrations of Triton-X 100 and sodium deoxycholate (SD), enhanced solubilisation of the cytoplasmic cellular and nuclear content. A milder surfactant, SD has shown enhanced preservation of gross microarchitecture and core composition of native ECM in soft tissue and basement membrane decellularisation ^53,54^. Agglutination of DNA on the tissue surface, often an issue with use of SD was addressed with addition of DNase I in the buffer ^7,55^. The enzyme – detergent extraction system ensured adequate removal of nuclear debris. Use of a recombinant nuclease in the protocol, reduced the risk of prion disease transmission seen with bovine sourced enzymes^56^.

Periodic washes with PBS and deionised water facilitated removal of cellular debris and detergent remnants. There is always a certain degree of ECM damage associated with any decellularisation process. In addition to the physical, chemical, or enzymatic disruptive methods that affect ECM composition and organisation, factors like spontaneous tissue breakdown and the release of intracellular proteases during decellularisation, unintentionally cause more damage ^7,57^. From a protocol optimisation perspective, our use of mild denaturing and processing conditions at 4°C at a neutral pH range of 7 – 8, for most of the duration, helped mitigate the impact of the decellularisation process on native pleural ECM.

Residual DNA in dPPM was less than 50 ng/mg of tissue, with a visible reduction in nuclei, reflecting protocol adherence to the criteria laid down by Crapo and Gilbert ^7,53^, for efficient decellularisation. Commercially available decellularised matrices derived from porcine SIS ^3^, porcine urinary bladder ^38^, and bovine dermis ^58^ bearing trace amounts of residual DNA have sought clinical applications in soft tissue reinforcement. Studies have looked at significant reduction in cellularity over complete cell removal as a balanced approach in tailoring tissue-specific decellularisation protocols^53,56,59^. This is mainly in consideration of preserving the core ECM structure and physiology for intended clinical applications.

Pleural ECM rich in collagen and elastin, confers viscoelastic characteristics, required in limiting lung distensibility during deep inspiration and generating elastic recoil pressure under resting conditions ^60^. Ultrastructural studies attribute the distinct organisational pattern of collagen and elastin fibres in an irregular interwoven plaited structure to be responsible for the mechanical resilience of lungs during respiration ^61,62^. The orientation and composition of the collagenous fibrous matrix equips the pleura to the physiological demands of flexibility along a multidirectional axis, a characteristic to lung volume changes during breathing.

Our choice of using freeze-thaw cycles in combination with low concentrations of milder surfactants, SD, and Triton X - 100 was in effort to ensure adequate removal of cells whilst retaining the biochemical and ultrastructural makeup of native PPM. dPPM characterisation however, reflected a degree of compromise to the native biochemical composition and microstructural integrity. Collagen and elastin histology exhibited decreased staining intensity, consistent with the reduction in insoluble collagen content quantified in dPPM. Microstructural changes to the dense irregular meshwork, characteristic of native pleural ECM, was evident with confocal imaging. Apparent changes in the angular distribution of fibre orientations in dPPM and the significance in the degree of this change, defined as coherency, may be a cumulative result of reduced collagen and elastin density with possible conformational changes in their core protein structure due to the denaturing effects of the decellularising agents used ^47,63,64^. These observations, however, require further investigations, more so as there is a considerable body of evidence associating fibre alignment and anisotropy with mechanical strength of bioscaffolds^65–67^ and functional behaviour therein.

Several studies have highlighted ultrastructural changes in ECM architecture using scanning electron microscopy and second harmonic generation ^64,68–70^. However, the conflicting degrees of ECM disruption for the same process in different tissues ^7,71,72^, makes it difficult to pinpoint the mechanism causing damage at varying levels during the decellularisation process. Research efforts by Hwang et al to trace spatial and temporal damage to collagen structure at a molecular level with the use of hybridising peptides ^63^, could be a solution in optimising tissue-specific decellularisation strategies, based on the molecular assessments of structural ECM damage with the routinely used decellularising agents like SDS, Triton – X or SD.

Increased membrane thickness and changes in ECM microarchitecture, did not severely alter the innate mechanical integrity of native PPM^73^. dPPM under uniaxial tension at a low strain rate, exhibited similar trends in viscoelastic deformation with linear and failure regions of response. This was reflected in the comparable Youngs modulus and tensile strength of dPPM, indicative of preserved native ECM stiffness and strength, respectively. Native membrane stiffness retained in the dPPM, is a key physical cue^74^ with potential to direct microscale cellular attachment, migration and subsequent cell-induced tissue remodelling at a macroscale level. The conserved mechanical strength in the derived dPPM could be of interest as an orthotopic TE graft in reinforcing sealing techniques and pleural repair.

The hydrogel component of pleural ECM is rich in PG and glycoproteins that function as mechanical buffers providing hydration and a repository of bound growth factors that modulate cell behaviour and tissue response to injury and wound healing. sGAG decorating the protein core of PG bind secretory factors including VEGF, EGF, TGF- β, and platelet derived growth factors to be incorporated within the ECM^36,72,75^. Significant reduction in sGAG content in dPPM is consistent with findings in other decellularisation studies^72,74,76^. Water-soluble sGAG are less resistant to decellularisation and are subsequently lost during repeated wash cycles. Although known for providing intrinsic compressive strength, reports suggest reductions in sGAG do not have an adverse effect on the gross mechanical characteristics of ECM. From a processing perspective, loss in sGAG improves diffusion of the decellularisation buffer and subsequent washes, enhancing removal of cellular debris ^52^. Additional processing of xenogeneic decellularised ECM is important in ensuring safety, biocompatibility, and off-the-shelf clinical potential. We assessed sterilisation efficiency of several known agents determining 0.1% peracetic acid to be most effective for dPPM. Routinely used as a mild decellularising agent with low cytotoxicity, PAA is known to be less disruptive to the mechanical and biochemical integrity of the ECM ^77,78^.

To maintain tissue relevance, we utilised a human pleural mesothelial cell line (MeT-5A) ^79^ to assess biocompatibility of dPPM in sustaining cell attachment, viability, and proliferation. Live-dead staining of the cells exposed to dPPM for a week, evidenced negligible cytotoxicity. Labelled MeT-5A cells seeded on sterile dPPM exhibited adhesion and proliferation. From a protocol optimisation objective, cytocompatibility of our dPPM scaffolds confirms minimal retention of decellularising and processing agents, (detergents, nucleases and sterilant) in the matrix, highlighting protocol safety and efficiency. Although decellularised porcine pleura is relatively unexplored, our study draws parallels with clinically approved bioprosthetic meshes in wound healing, such as porcine urinary bladder matrix (MatriStem®)^7^ and porcine mesothelial grafts (Meso Biomatrix®) ^80^, representing a step forward towards developing pleural acellular scaffolds for applications in lung tissue repair and regeneration.

## Conclusion

Our methodology for decellularising porcine pleura using freeze-thaw cycles and low concentration detergent-enzyme buffer, ensured adequate cell removal while preserving gross biochemical and mechanical attributes of native pleural ECM. Nuclear staining and residual DNA content in dPPM met the essential criteria for effective cell removal. Although biochemical composition and structure reflected the anticipated changes in native ECM, the mechanical strength and behaviour of dPPM was comparable with native pleural tissue. Sterilised dPPM scaffolds facilitated cellular attachment, viability, and proliferation with negligible cytotoxicity. To summarise, our study lays the groundwork in exploring decellularised porcine pleurae as site-specific acellular surrogates in lung tissue engineering applications.

## Acknowledgements

This research was funded by the Engineering and Physical Sciences Research Council (EPSRC) Centre for Doctoral Training in Regenerative Medicine (EPSRC reference EP/L015072/1) and a project grant from the North Staffordshire Medical Institute (Stoke - on – Trent, UK).

## Author contributions

**Trisha Vikranth**: Investigation, methodology, formal analysis, visualisation, writing – original draft.

**Dr Tina Dale**: Supervision, project administration.

**Professor Nicholas Forsyth**: Conceptualisation, funding acquisition, supervision, writing – review and editing.

## Ethics approval and consent to participate

Our study did not require ethical approval as the porcine tissues were obtained via food chain supply and not through a veterinary or research-linked facility.

## Notes

### Competing Interest Statement

The authors have declared no competing interest.

